# Experimental Autoimmune Encephalomyelitis Causes Skeletal Muscle Dysfunction in Mice

**DOI:** 10.1101/2025.03.04.641397

**Authors:** Julian Boesch, Pamela Ramseier, Sarah Tisserand, Eliane Pierrel, Giuseppe Locatelli, Serge Summermatter

**Author notes:** Corresponding author: Serge Summermatter. Contributed equally (Julian Boesch, Pamela Ramseier).

## Abstract

Multiple sclerosis (MS) is a neuroinflammatory disease affecting the brain and spinal cord and characterized by demyelination, neurodegeneration and chronic inflammation. More than 90% of people with MS present with peripheral muscle dysfunction and a progressive decline in mobility. Current treatments attenuate the inflammatory processes but do not prevent disease progression. Therefore, there remains an unmet medical need for new and/or additional therapeutic approaches that specifically improve muscle function in this patient population.

The development of novel treatments targeting skeletal muscle dysfunction in MS will depend on suitable preclinical models that can mimic the human musculoskeletal manifestations of MS. Using a non-invasive approach to assess muscle function, we demonstrate *in vivo* that Experimental Autoimmune Encephalomyelitis (EAE) impairs skeletal muscle strength. Our data reveal a 28.3% (*p*<0.0001) lower muscle force in animals with EAE compared to healthy control mice during electrically evoked tetanic muscle contractions that occur at intervals of 0.25 seconds and thus mimic fatiguing tasks.

As we conduct force measurements by direct transcutaneous muscle stimulation in anesthetized animals, our setup allows for the repeated evaluation of muscle function, and in the absence of primary fatigue or reduced nerve input which constitute important confounding factors in MS. Taken together, our data highlight important similarities between MS in humans and EAE in mice with regards to skeletal muscle contractile impairments, and provide first evidence for a non-invasive *in-vivo* setup that will enable the preclinical profiling of novel drug candidates directed at specifically improving muscle function in MS.

## INTRODUCTION

Multiple sclerosis (MS) is a chronic inflammatory disease characterized by immune cell infiltration into the central nervous system (CNS), diffuse glial activation, axonal demyelination and neurodegeneration. Common symptoms of MS include fatigue and mobility impairments, which are reported by 37–78% and >90% of people with MS (pwMS), respectively (Hemmett et al., 2004; Oliva Ramirez et al., 2021; Locatelli et al., 2024).

Fatigue is defined as a “subjective sensation of weariness, an increasing sense of effort, a mismatch between effort expended and actual performance, or exhaustion” (Kluger et al., 2013). This condition can result from CNS damage (primary fatigue) or only indirectly be related to MS (secondary fatigue) (Patejdl and Zettl, 2022). Secondary fatigue can be due to sleep disturbances, chronic urinary tract infections, side effects of pharmacological interventions, or impairments at the musculoskeletal level (Patejdl and Zettl, 2022). The latter is referred to as motor fatigability and describes the reduced capacity of skeletal muscle to produce and maintain voluntary or evoked force during physical activity (Patejdl and Zettl, 2022; Royer et al., 2022). It has been shown that motor fatigability remains more pronounced in pwMS than in healthy controls even when evoking contractions directly on muscle by electrical stimulation, thus bypassing the CNS and eliminating the contribution of primary fatigue (de Haan et al., 2000). These data suggest that motor fatigability in MS is not inextricably linked with primary fatigue. Instead, muscle-intrinsic alterations drive or at least importantly contribute to motor fatigability in MS.

Indeed, detailed investigations into human skeletal muscle morphology revealed mild atrophies of the quadriceps (rectus femoris) and the gastrocnemius muscle in pwMS (Kirmaci Zİ et al., 2022). These atrophies pertain to specific fiber types, particularly to type IIA fibers (Hansen et al., 2015). Noteworthy, type IIA fibers contribute to both fatigue resistance and muscle strength and are recruited for tasks requiring greater muscle strength and fatigue resistance (Hodson-Tole and Wakeling, 2009). In addition, impairments in the metabolic capacity have been found in muscle tissue of pwMS. To be precise, analyses of mitochondrial activity in skeletal muscle showed a reduced expression of complex I and II of oxidative phosphorylation (OXPHOS) (Locatelli et al., 2024).

Importantly, current treatments for MS are not directed at restoring any of these musculoskeletal impairments. Rather, these treatments, including interferons, glatiramer acetate, teriflunomide, sphingosine 1-phosphate receptor modulators, fumarates, cladribine, and monoclonal antibodies targeting either CD20, integrins, or CD52, were designed to induce anti-inflammatory effects systemically and/or in the CNS (McGinley et al., 2021). In addition, there is a potassium-channel blocker that is prescribed to treat walking disabilities in pwMS. This inhibitor is called Fampridine and improves gait balance in subjects with MS (Ghorbanpour et al., 2023). Fampridine acts on the central and peripheral nervous systems and is the only pharmacological therapy that has been approved for gait imbalance in these patients (Ghorbanpour et al., 2023). Overall, given the limited treatment options for musculoskeletal dysfunctions in MS, there is an unmet medical need for adjunct therapies to restore functional independence.

Here we investigated whether Experimental Autoimmune Encephalomyelitis (EAE) as a rodent model for MS preclinically mimics key musculoskeletal impairments that occur in pwMS, and whether these impairments can be monitored non-invasively *in vivo*.

## MATERIALS AND METHODS

### Animals

All animal studies described were performed according to the official regulations effective in the Canton of Basel-City, Switzerland. The mice were housed at 25 °C with a 12:12 h light-dark cycle and fed a standard laboratory diet (Nafag, product # 3890, Kliba, Basel, Switzerland). Food and water were provided *ad libitum*. Prior to the study the animals were acclimatized to the research facility in Basel (Switzerland) for 7 days. EAE was induced as previously described (Ivan et al., 2023). In brief, 8 weeks old, female C57Bl/6 mice were subcutaneously injected with 200µg rat Myelin Oligodendrocyte Glycoprotein (MOG) emulsified with 4mg/mL complete Freund’s adjuvant and intraperitoneally injected with 100ng pertussis toxin. Two days later the animals received a pertussis toxin boost. Body weight was continuously monitored throughout the study and the clinical disease course was assessed using a 0–4 score scale. Mice displaying a 3–5% weight loss but no other motor impairments received a score of 0 (weight-loss stage), animals showing a limp tail were scored as 0.5-1 (day of clinical onset), mice displaying a partial weakening of hind limbs received a score of 1.5-2 and animals presenting hind limb paraparesis/paraplegia were scored with 2.5-3 (symptomatic disease peak). Mice displaying hindlimb paralysis and forelimbs paraparesis were scored with 4 and met termination criteria. On day 31 post disease induction, muscle fatigability of the left leg was measured. On day 32, the animals were euthanized, and muscle tissues were collected from the right, unstimulated leg.

### Neurofilament Light Chain (NF-L), Glial Fibrillary Acidic Protein (GFAP) and Insulin-like Growth Factor 1 (IGF-1) ELISA

GFAP (Creative Diagnostics – Ref#DEIA7378), NF-L (Uman Diagnostics – Ref#10-7001) and IGF-1 (Rockland – Ref#KOA0195) ELISA were performed in cerebrospinal fluid (CSF) diluted 1/50 and 1/125, and in plasma diluted 1/5 in the sample diluent provided in each kit, respectively. Diluted sample and standard (in duplicates) were incubated on the coated plate, followed by incubation with the biotinylated antibody. After several washing steps, the HRP-conjugated antibody was added and the amount of GFAP or NF-L was revealed by a TMB (tretramethylbenzine) solution. The reaction was stopped by addition of the stop solution, and the plate was measured at 450nm in a Spectramax340 photometer.

For the analysis, the optic density of the blank was subtracted from each measurement. The concentrations of NF-L, GFAP and IGF-1 in the well were extrapolated from the linear regression. Finally, the dilution factor was applied to determine the sample concentration.

### Muscle force measurement

Motor function of the left hind leg was measured non-invasively using a setup described previously (Boesch et al., 2022). In brief, the animals were anesthetized and electrodes for transcutaneous stimulation were put in place on the shin and the thigh. The foot was then positioned on a homemade pedal connected to a force transducer. Muscle contractions were evoked via electrical stimulation of the hind leg through the transcutaneous electrodes and the force generated was recorded. In contrast to the method previously described, stimulation of the leg muscles was done at a low frequency (40 Hz) with a new tetanic stimulus every 0.25 seconds and the force generated was recorded for 120 stimulations to evaluate motor fatigability.

### Gene expression profiling

Total RNA was extracted from skeletal muscle using TRIzol reagent (Invitrogen). Reverse transcription was performed with random hexamers on 1 μg of total RNA using a high-capacity reverse transcription kit (Applied Biosystems), and the reaction mixture was diluted 20-fold. RT-PCRs were performed in duplicates in 384-well plates on an AB7900HT cycler (Applied Biosystems) using specific TaqMan probes (Applied Biosystems). Data were normalized to two housekeeping genes using the ΔΔCT threshold cycle (CT) method. Fluorescence was measured at the end of each cycle, and after 40 reaction cycles, a profile of fluorescence versus cycle number was obtained. Automatic settings were used to determine the CT. The comparative method using 2−ΔΔCT was applied to determine the relative expression. Results are expressed as fold changes over controls.

### Statistical analyses

Statistical analyses were performed using *t*-tests in Prism 10 (GraphPad Software, Inc., La Jolla, CA). As the sole deviation, analyses of CSF and plasma samples were conducted using a nonparametric test (Mann-Whitney U) in Prism 10 (GraphPad Software, Inc., La Jolla, CA. Differences were considered to be significant when the probability value was <0.05.

## RESULTS

EAE overall exerted limited lasting effects on body weight in mice (Figure 1 a). Following disease induction, the animals showed an initial drop in body weight (probably due to the stress associated with the injections) on the first day, but then readily resumed normal growth. Around day 12, the body weight started to decline rapidly (Figure 1 a and b). The average decrease in body weight reached 8% maximally, was only transient and coincided with the onset of EAE (Figure 1 c). In fact, the animals already caught up growth again as of the 16^th^ day post disease induction (Figure 1 b). In accordance, the clinical scores accumulated until day 16. Thereafter, a remission phase set in (Figure 1 d).

**Figure 1:**
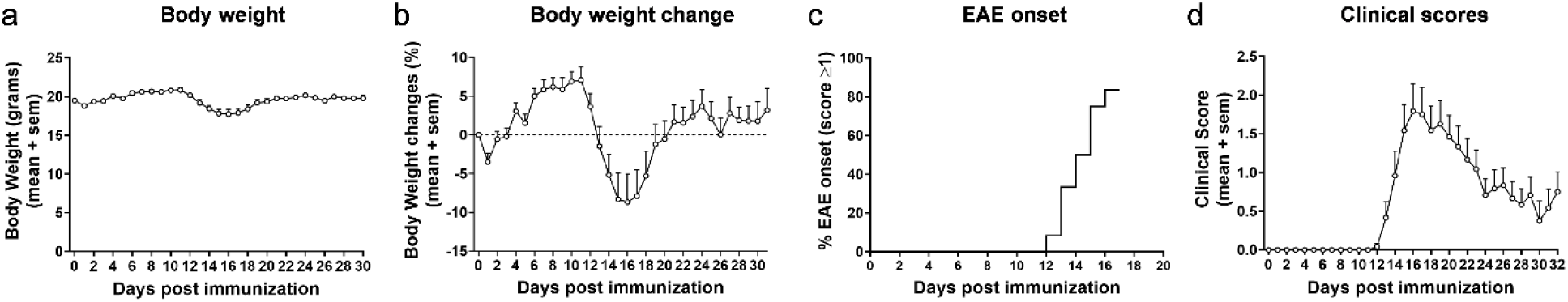
Clinical parameters in EAE. Body weight (a) and body weight change (b) in female mice following experimentally induced autoimmune encephalomyelitis. Percentage of animals showing first signs of EAE in relation to the timepoint of symptom onset (c) and clinical scores (d). Data in a-d have only been collected for mice with EAE (not for sham animals). n=12. Values are expressed as mean ± SEM.

The EAE model is a model of relapsing-remitting MS and is thus characterized by alternating phases of acute flare-ups and symptom remissions. To avoid confounding caused by acute inflammation during a flare-up, all *in-vivo* and *ex-vivo* measurements were conducted following symptom remission. More precisely, skeletal muscle function was assessed on day 30, when the animals had returned to their normal body weight and clinical symptoms had partially subsided (Figure 1 b and d). However, the disease was active at this time point, as indicated by the significant upregulation of NF-L, a biomarker of neuronal injury, in the CSF (Figure 2 a). Moreover, elevated levels of GFAP were detectable in the CSF, hinting towards persisting astrocyte activation and astrogliosis (Figure 2 b). By contrast, IGF-1 as a circulating growth hormone altering muscle mass was similar between the two groups (Figure 2 c). Using repeated tetanic muscle stimulations *in vivo*, we found that animals with EAE displayed lower muscle strength during a protocol that simulated fatiguing tasks than their healthy controls (Figure 3 a). Quantification of the areas under the curve further highlighted a significant overall reduction of -28.3% (*p*<0.0001) in muscle strength in mice with EAE (Figure 3 b).

**Figure 2:**
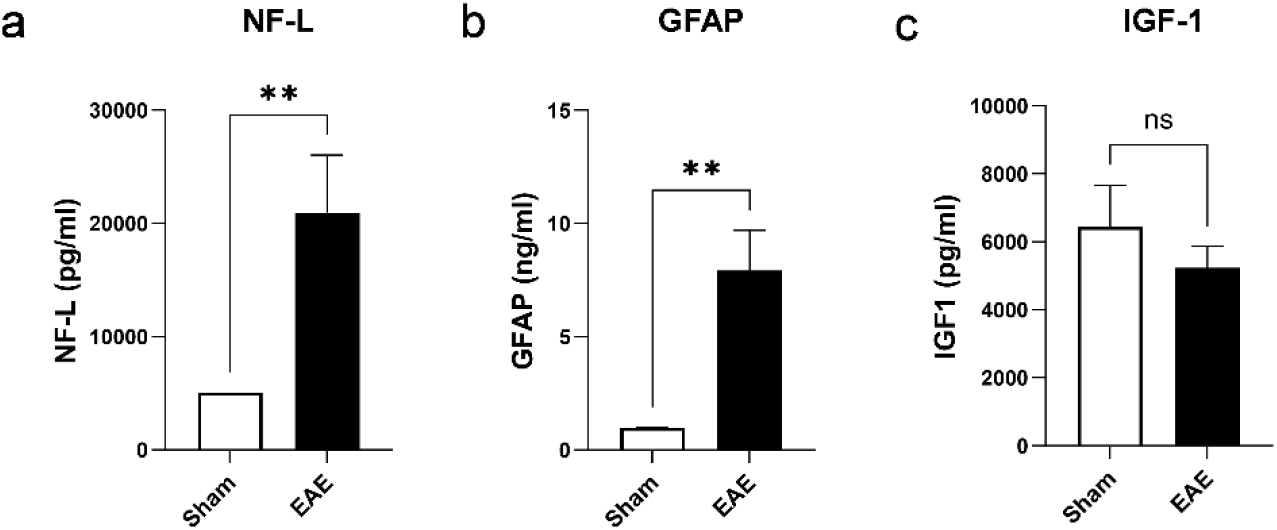
Biomarkers of axonal damage, neuroinflammation and muscle growth. Levels of NF-L as biomarker of axonal damage (a) and GFAP as biomarker of astrocyte activation and astrogliosis (b) in cerebrospinal fluid. Plasma levels of IGF-1 as growth hormone regulating muscle function (c). n=4 for sham and n=12 for EAE. Values are expressed as mean ± SEM.

**Figure 3:**
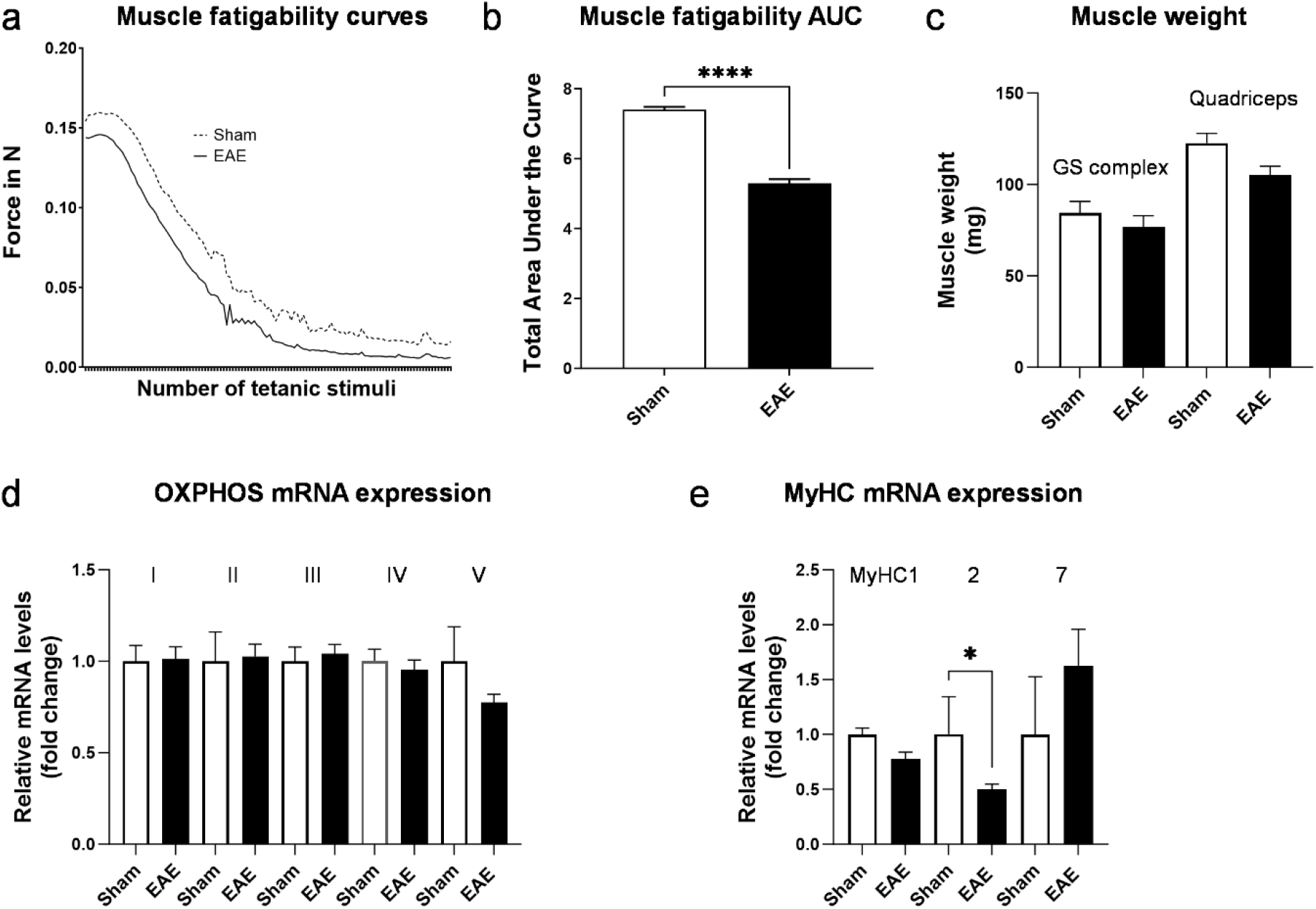
Characterization of musculoskeletal function in EAE. Excursion curves of muscle strength during fatiguing muscle contractions induced by transcutaneous electrical stimulation (a) and corresponding area under the curve (AUC) (b). Muscle weight of the gastrocnemius-soleus complex (GS) and quadriceps muscle (c). Relative mRNA expression of OXPHOS subunits; Ndufb5 for complex I, SDHB for complex II, Uqcrc2 for complex III, Cox5b for complex IV and ATP5o for complex V (e). Relative mRNA expression of myosin heavy chains (e). n=4 for sham and n=10-12 for EAE. Values are expressed as mean ± SEM.

To evaluate whether similar changes in muscle weight occur in EAE and MS in humans, we determined in this study the muscle weight and gene expression of myosin heavy chains (MyHC) and OXPHOS components *ex vivo* (post-mortem). Muscle weight of larger muscle groups, such as the gastrocnemius-soleus complex and the quadriceps muscle were not statistically significantly different between groups, although the quadriceps showed a trend towards lower muscle weight in diseased mice (−14.3%, *p*=0.06) (Figure 3 c). A reduced metabolic capacity and altered contractile properties have been reported in skeletal muscles of pwMS (Locatelli et al., 2024). Therefore, we assessed the mRNA expression of representative subunits of the five complexes that mediate OXPHOS. No differences in the mRNA expression of OXPHOS components in skeletal muscle were detected between the two groups (Figure 3 d). However, we observed a significant reduction by 50.2% (*p*>0.05) in MyHC2 mRNA expression in the muscles of mice with EAE (Figure 3 e). MyHC2 is predominantly expressed in type IIA muscle fibers. In contrast, the expression of MyHC1, which is found in IIX/D fibers, and the expression of MyHC7, which prevails in type I fibers, were not differentially expressed between sham or animals with EAE. Intriguingly, specific atrophies of IIA fibers have been reported in pwMS (Hansen et al., 2015). As these fibers are important to generate and maintain forces over a prolonged period, the reduced expression of MyHC2 in the EAE model is consistent with the observed functional deficits.

## DISCUSSION

Mobility disability is an important contributor to disease burden in pwMS (Hemmett et al., 2004; Locatelli et al., 2024). Treatment options to address the underlying muscle dysfunctions are limited and there is a high unmet medical need for treatments that can complement current therapies. To develop such novel treatments, preclinical models for specific features of MS are required. Those animal models need to reflect human disease patterns to have the potential for clinical translatability. The EAE model is a widely-used animal model for MS, but its impact on the musculoskeletal system has not been well-characterized.

While attempts have been made to investigate muscle function in EAE, these studies suffer from a couple of limitations. For instance, they either relied on *ex-vivo* analyses of isolated muscles not yet shown to be heavily affected by MS in humans (e.g., soleus, extensor digitorum longus etc.), or used invasive *in-situ* procedures (de Haan et al., 2004; Spaas et al., 2021). A further disadvantage of those methods is that they are terminal and do not allow for continuous monitoring of muscle strength. To address such limitations, we established a setup that is completely non-invasive, measures the function of larger muscle groups critical for mobility, and can be applied repeatedly on individual animals. We have shown here that in an experimental setup which mimics fatiguing tasks, muscle strength is significantly reduced in animals with active EAE compared to healthy controls. These differences could be observed in the chronic stage of the model, in which acute inflammation has subsided but neurodegeneration/neuroinflammation remain detectable in the mouse CNS as evidenced by MS-relevant biomarkers such as GFAP and NF-L in the CSF. Notably, at the same stage, systemic factors potentially influencing muscle mass such as IGF-1 were not different from control mice. Thus, our method enables the assessment of baseline values prior to profiling a novel drug candidate and the subsequent evaluation of the efficacy of specific drug substances directed at altering muscle function. The suitability of the EAE model is further underscored by the fact that the model displays alterations in MyHC expression that are reminiscent of the MyHC changes reported in pwMS. Our findings hold promise for the future development of novel drug candidates improving muscle function and restoring mobility in pwMS.

## ACKNOWLEDGEMENTS

We thank Steven Kovacs for his valuable comments on the manuscript.

